# Working memory affects motor, but not perceptual timing

**DOI:** 10.1101/2024.06.25.600202

**Authors:** MohammadAmin Farajzadeh, Mehdi Sanayei

**Affiliations:** School of Cognitive Sciences, Institute for Research in Fundamental Sciences (IPM), Tehran, Iran; Shahid Beheshti University of Medical Sciences, School of Medicine, Tehran, Iran

**Keywords:** Time perception, motor timing, perceptual timing, working memory

## Abstract

Whether different timing tasks utilize the same brain processes is still debated. To address this question, we investigated how working memory affects two different timing tasks: time reproduction and time discrimination. We found that delay intervals led to an overestimation in the reproduction task but did not introduce any bias in the perception of time in the discrimination task. Delay intervals affected the perception of time when subjects had to actively reproduce the perceived interval, but not when subjects were merely recalling the content of working memory. In subsequent Bayesian modeling, we showed that in the reproduction task, subjects updated their measurement of the stimulus on the current trial (likelihood) based on the delay interval, rather than changes in the motor system or updating priors based on the delay interval. Our findings suggest that the brain processes involved in time reproduction and discrimination are not completely overlapping, and that delay intervals in working memory tasks lead to changes in updating the likelihood. This robustness in prior and updates in likelihood provides both stability and sensitivity in the perception of time.

## Introduction

Perception of time and duration of events is one of the most essential parts of everyday life and have a pivotal role in speech processing, decision making and motor control ^1–3^. Perception of sensory information (including time) and keeping this information in memory are considered different processes, as studies reveal distinct brain circuits for perception vs. memory ^4^. However, the capacity to perceive time and different durations is intertwined with the working memory process ^5–9^. The classic psychological model for time perception, which is still in use today, consists of a pace-maker and an accumulator, in which the pace-maker emits pulses and these pulses are accumulated to be compared with the reference duration in working memory^5,10^. Despite the importance of the role of working memory in time perception, this topic has remained relatively under-studied (but see references 11 and 12).

In time perception literature, there are two main tasks used to study time perception. Perceptual timing tasks (e.g., time discrimination), in which the subject is presented with two intervals and has to report which interval is shorter or longer ^11^. Subject does not actively reproduce either the first or second duration. On the other hand, there are motor timing tasks (e.g., time reproduction), which demands active engagement and manipulation of the perceived duration. In reproduction tasks, the experimenter presents a time interval to the subject, and the subject is asked to reproduce the presented time interval (e.g., by pressing a button for the perceived duration).

Working memory involves memorizing some information for short time periods ^12^. Working memory has been studied using tasks that require delayed responses to stimuli presented earlier^4,13^. Various studies have demonstrated that as this delay interval increases, the subject’s performance decreases^14^. Additionally, it has been shown that the presentation of certain types of distractor stimuli during the delay period leads to a larger decrease in working memory performance ^15^.

We can use delayed-response tasks to study the effect of working memory on time perception, but the delay interval itself is an elapsed time. So, the delay interval has dual effects on the perception of time. First, delay affects the task as a component of any working memory task. Second, given the sample interval and delay interval are both time intervals, delay intervals can interfere with the perception of the sample interval, like a serial dependence effect ^16,17^. Serial dependence effect is a phenomenon in which stimuli from previous trials could have an effect on the perception of the current trial (i.e., local context^18^). Recent studies showed that this local context effect in time perception could be due to the strong effects of previous trials (updating prior). On the other hand, local context could be due to effects of previous trials on the signal to noise ratio of the stimulus on the current trial (updating likelihood). Although these studies investigated the effect of previous trials, none of them studied the effect of the delay interval, which comes after the to be timed interval, on time perception, either with a reproduction or a discrimination task.

In this study, we examined the effect of delay interval on time perception using both reproduction and discrimination tasks in the same subjects. We found that longer delay intervals led to an overestimation of the sample interval in the reproduction but not in discrimination task. We showed that, in the reproduction task, time perception attracted towards the delay interval duration. Furthermore, we proposed a computational model to explain this overestimation of sample interval. We found stronger evidence that the delay interval updates the likelihood function rather than the prior or changing the motor noise.

## Material and Methods

### Subjects

Fourteen subjects (5 female, 25.5 ± 5.16 years old) participated in this study. All subjects have normal or corrected-to-normal vision, and were naïve about the goal of the experiment, except the two authors (subject number 1 & 2). All subjects were enrolled in both reproduction and discrimination tasks and the order of the tasks was counterbalanced between them. This study was conducted under the approval of the ethics committee at the School of Cognitive Sciences, Institute for Research in Fundamental Sciences (IPM). A written informed consent was taken from subjects prior to the experiment.

### Apparatus

Experiments were performed on a PC with Linux operating system. We used the Psychtoolbox 3 extension with MATLAB (2016b) for designing our tasks ^19–21^. Stimuli were presented on a 17” monitor (refresh rate 60Hz) placed ∼60cm from subjects. Subjects sat on a chair in a dim room, and their head was stabilized with a chin and head rest. The position of the left eye was monitored (1KHz, EyeLink 1000, SR Research, Mississauga, Ontario) during the experiment.

### Time reproduction task

On each trial, we presented a fixation dot (0.2°) and two targets (left target: 0.5° diameters, right target: 2° diameter, 10° eccentricity) to the subject (Fig. 1A). After the subject fixed the central dot for 200ms (within 2° window), a circular wheel-shaped stimulus (diameter = 2.5°) was shown. The duration of the stimulus was 400, 600 or 1000ms, chosen pseudo-randomly on each trial. After that, there was a delay in which only the fixation point and targets were present. The delay was selected pseudo-randomly on each trial from 500, 900, or 1700ms. After this delay, we presented a second stimulus, which was the same as the first one but rotated by 180°. The subject had to make a saccadic eye movement to the right target when the second stimulus’ duration reached the first stimulus’ duration. The reproduced time was measured from the onset of the second stimulus on the screen until the subject made the saccade. After response, the difference between the duration of the presented stimulus and the reproduced time was presented to the subject with millisecond resolution at the location of the right target. Each participant completed 9 blocks, and each block consisted of 72 trials. Before the main reproduction task, each subject did 4 blocks (18 trials each, 2 repetitions per condition) of the reproduction task, to get them familiarized with the task. The trials were excluded from any analysis.

**Figure 1.**
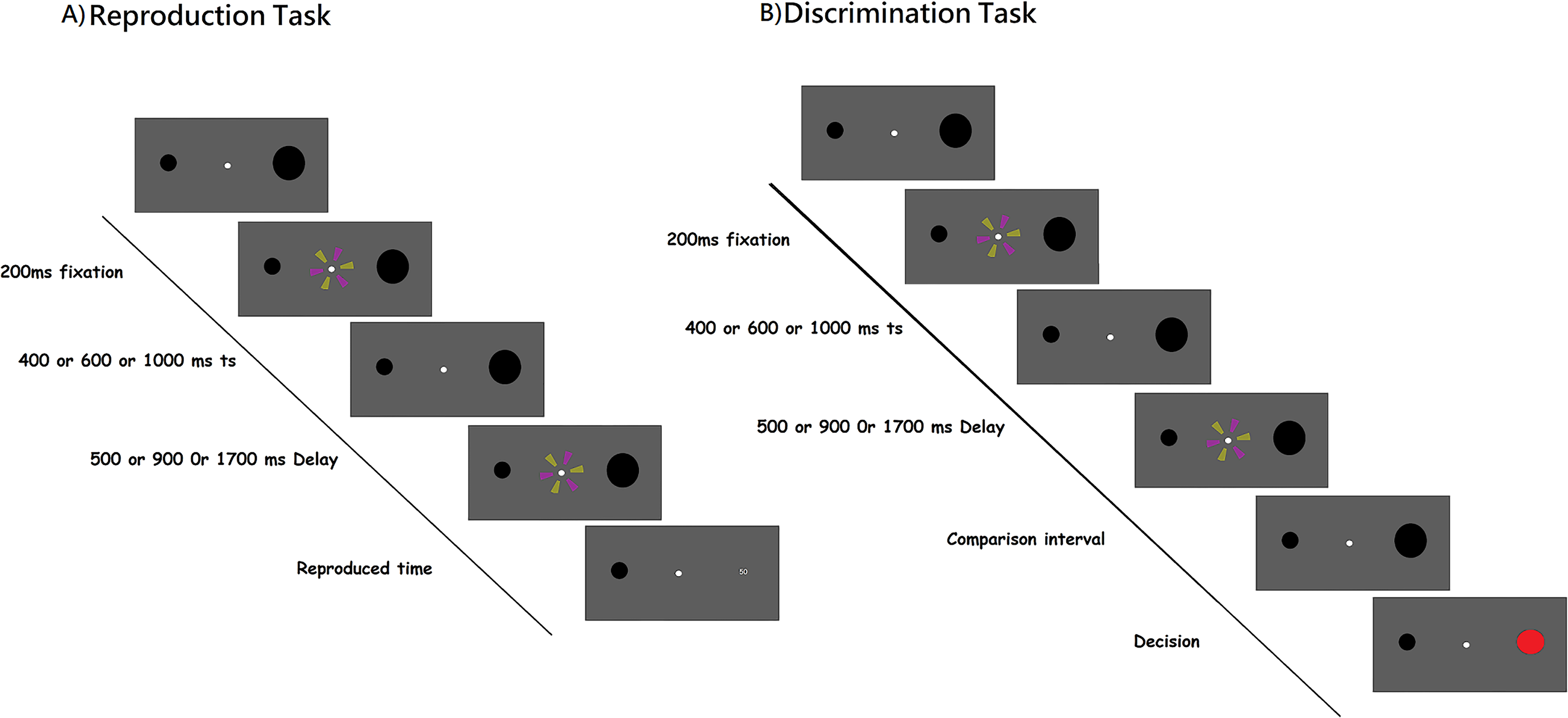
The sequence of events in a trial for reproduction (A) and discrimination (B) tasks. In the reproduction task (A), after the subject acquired fixation, a sample stimulus presented on the screen for the duration of t_s_. Then, the subject had to wait for a delay interval (t_d_). After that, they reproduced the t_s_ by making a saccade to the right target. At the end of each trial, we showed the response error (difference between the sample interval and reproduced duration) to the subject in millisecond. In the discrimination task (B), the subject was shown two intervals, with a delay (t_d_) between them and they had to compare them. The subject had to report which interval was longer with a saccadic eye movement to one of two targets. At the end of each trial visual feedback (correct or error) was presented to the subject.

### Time Discrimination task

In this task, we presented two stimuli sequentially with a delay between them and then asked subjects to indicate which one of the presented stimuli was longer (Fig. 1B). The design and timing of the discrimination task were similar to the reproduction task, with the exception of the duration of the second stimulus. After the delay interval, the second stimulus’ duration was the first stimulus’ duration ± 6, 12, 24, 48% of the first stimulus duration. The subject had a 4-second window to respond. If the second stimulus lasted longer than the first stimulus, the subject had to make a saccade to the right target. If the second stimulus lasted shorter than the first stimulus, the subject had to make a saccade to the left target. After that, the feedback was provided for the subjects with red and green filled circles, for correct and incorrect responses, respectively. Each participant carried out 18 blocks, and each block consisted of 72 trials. Before the start of this task, subjects did blocks of 18 trials, with only the easiest conditions. After a subject performed 100% on two consecutive blocks, we moved on to the main task. On average, subjects performed 4.35 ± 2.6 (mean and standard deviation) blocks to get familiarized with the task. These trials were not included for further analysis.

### Analysis of behavioral data

For the reproduction task, we excluded any reproduced time that was larger than 1.5 IQR for each subject (0.7% of trials from all subjects excluded) ^22^. For the remaining trials, we calculated the mean and standard deviation of the reproduced time for each sample duration and each delay interval. We plotted the mean reproduction time (𝑡_𝑟_) against sample interval duration (𝑡_𝑠_) for each subject and applied a linear regression function to the data from all subjects. These lines crossed the diagonal line at the ‘indifference point’, which represent an interval where subjects had a veridical perception of the physical time ^23^. We also calculated the indifference point for each subject.

For the discrimination task, for each combination of sample stimulus duration and delay interval (9 in total), we fitted a cumulative Gaussian function to the percentage of ’longer’ response as a function of the comparison stimulus duration. To assess how well the cumulative Gaussian fitted the data, we measured the goodness of fit and excluded poorly fitted conditions (r^2^ < 0.5) (none was excluded) ^22^. We measured the point of subjective equality (PSE) as the mean of the fitted Gaussian and its standard deviation as an index of variability for the discrimination task. Further, we fitted linear regressions to the PSE values for each delay interval as a function of sample interval duration to obtain indifference points the same as in the reproduction task. To investigate the effect of delay in each task on the indifference point, we applied a One-way repeated measure analysis of variance (RM-ANOVA) on each task, separately.

We applied a three-way RM-ANOVA to the mean of the reproduced time and PSE of the discrimination time to measure the effects of task (1 levels), sample interval (2 levels), and delay interval (3 levels) and their interaction on the accuracy of timing. We applied the same RM-ANOVA to standard deviation of both tasks to investigate the effects of these factors on the precision of timing. To investigate the effect of delay duration on the two tasks separately, we applied a two-way RM-ANOVA on the mean of time perception to measure the effect of sample interval duration and delay interval duration on the mean of the reproduced time (reproduction task), or the PSE (discrimination task).

To analyze the delay dependency of the reproduced interval, we used a deviation index. In the reproduction task, deviation index on each trial was computed as the reproduced duration minus the mean of reproduced durations for all trials with that sample interval. In the discrimination task, we first transformed data from the cumulative Gaussian format to a Gaussian distribution with the same mean and standard deviation, for each sample and delay interval, per subject. Then we draw 500 samples for each sample and delay interval combination (9 conditions in total).We then treated these distributions the same as the reproduced time, as previously has been done ^22,24^, and calculate the deviation index similar to how we calculated in the reproduction task.

Positive values for the deviation index showed that the reproduced interval for the current trial was longer than the mean reproduced duration and negative values indicated that the reproduced duration was shorter than the mean reproduced duration. To measure the effect of the delay duration on the deviation index, we fitted a simplified Derivative of Gaussian (DoG) curve to describe the deviation index as a function of difference between the current sample interval and delay duration as:

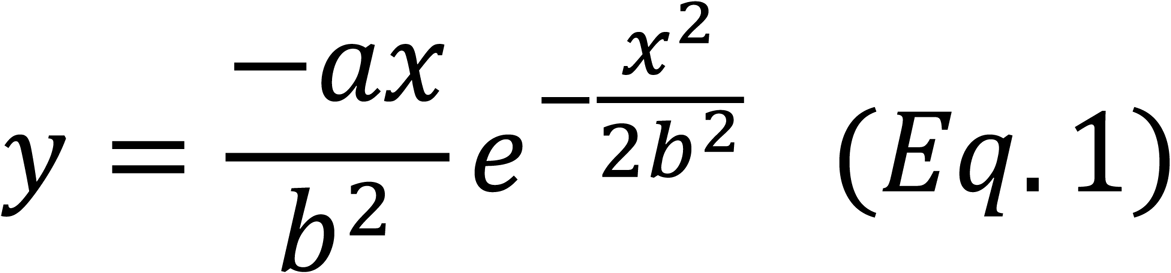

Where y is the deviation index, x is the difference between sample interval and delay interval, a and b are fitted parameters. We fitted DoG at both the group and individual levels. To measure the goodness-of-fit, we computed r^2^ for DoG model compared to non-model and linear interaction between deviation index and delay interval.

### Modeling

To model the effect of delay interval on the reproduced time, we used a combination of Bayesian Observer^25^, distractor ^26^, and mixed-Gaussian update models^18^ . The basic observer consists of three different components: likelihood function, prior distribution and motor noise. Subjects could update any of these components based on the delay interval on each trial. To investigate which of these three possibilities explained our data best, we used modified versions of the classic Bayesian model for time perception ^25^. We first explain the model in general form, and then the modifications that we applied for our three models.

This model breaks the process of time perception to measurement, estimation, and production. In the measurement step, the sample interval presented to the subject (𝑡_𝑠_), is mapped to measurement time, 𝑡_𝑚_, using a Gaussian distribution, where the mean of this Gaussian distribution is equal to 𝑡_𝑠_, and its standard deviation is equal to 𝑡_𝑠_ × 𝑤_𝑚_, where 𝑤_𝑚_ is a free parameter in the model. We denote this step with likelihood function 𝜆.

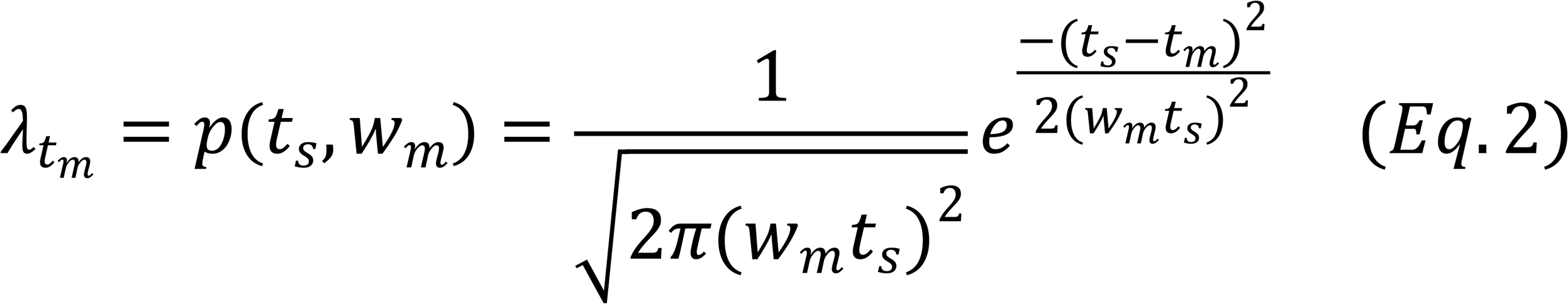

We defined the prior distribution of 𝑡_𝑠_ as a uniform distribution of times between 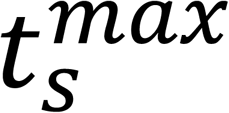 and 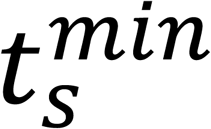 as follows:

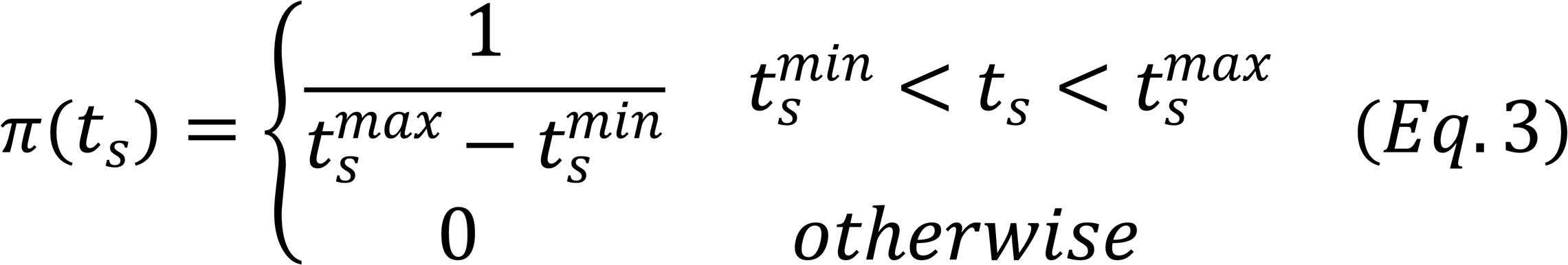

Posterior is the product of prior multiplied by likelihood function as:

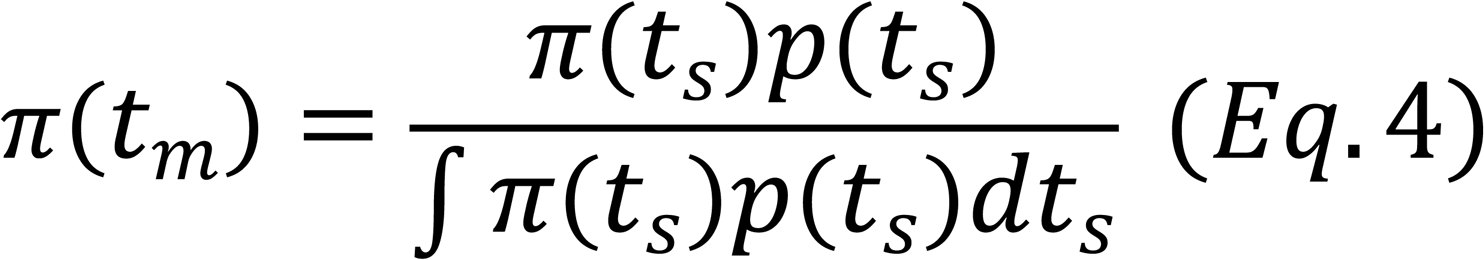

In the next step, which was estimation, the model combines this likelihood with the prior distribution of time samples based on the mean of the posterior, generating the estimated time (𝑡_𝑒_):

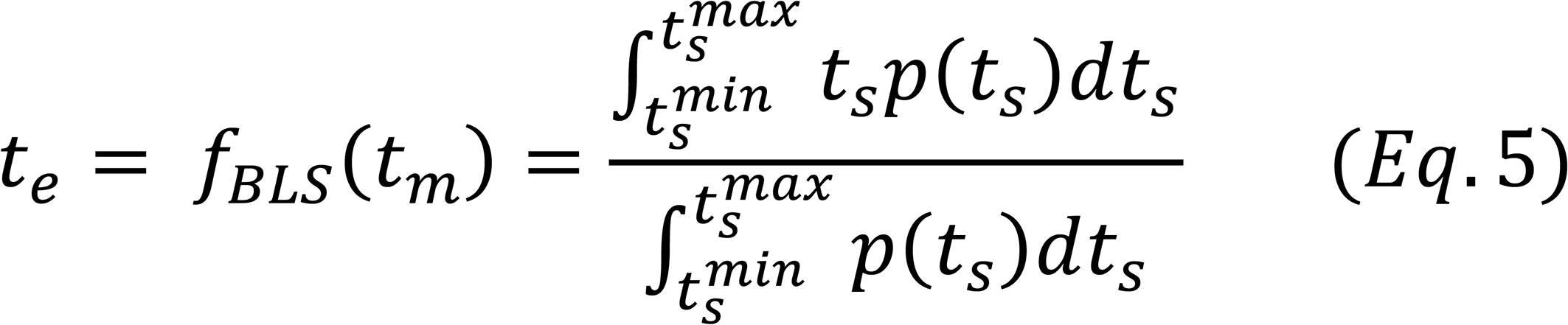

In the final step, the values of 𝑡_𝑒_ are mapped to the produced time (𝑡_𝑝_) using a Gaussian distribution, where the mean of the distribution is equal to 𝑡_𝑒_ and the standard deviation of the distribution is equal to 𝑡_𝑒_×𝑤_𝑝_, where 𝑤_𝑝_ is a free parameter in the model.

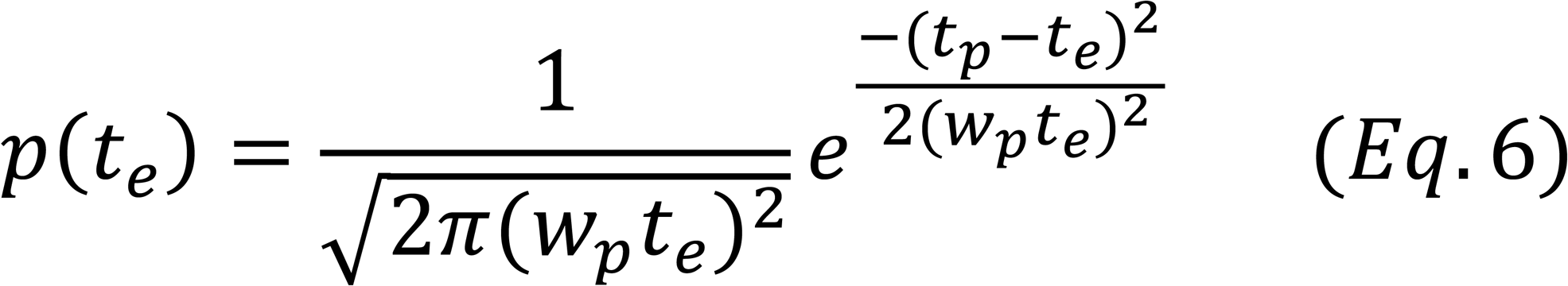

Finally, the direct relationship between the reproduced time and the sample time that was presented to the subject is explained by the following equation:

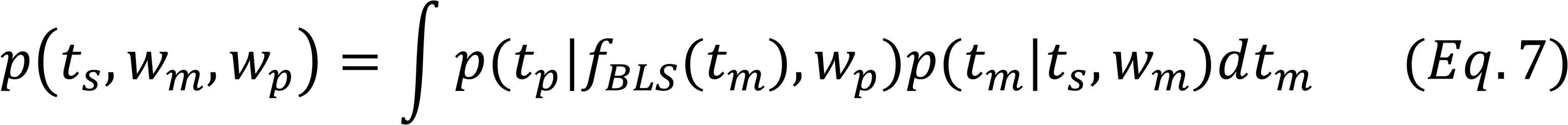

Now we are going to add component of distraction to model the effect of the delay duration (𝑡_𝑑_). The distraction effect on any duration, 𝑡, can be modeled by weighted average between 𝑡_𝑑_ and 𝑡:

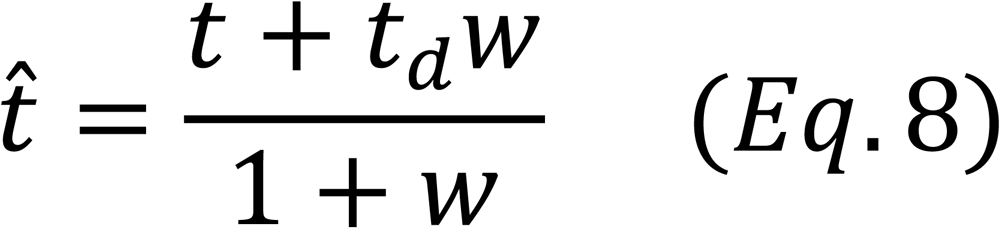

The weight of the distractor, 𝑤, was between 0 and 1. The greater the weight of the distractor, the more effect it has on the perceived duration (𝑡^). 𝑤 is computed as:

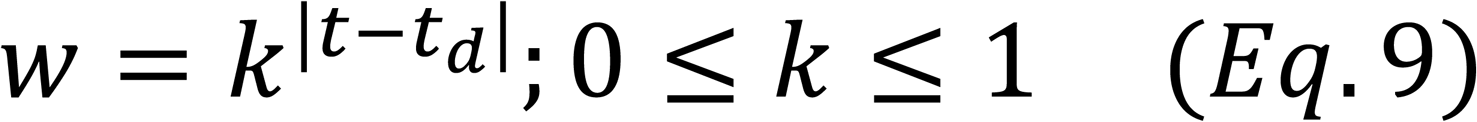

Where 𝑘 is the leak factor determining the distractor weight on the interval. When 𝑘 = 1 the weight will always be one. When it is any value between 0 and 1, as the difference between 𝑡_𝑑_and 𝑡 increases the weight and the effect of the distractor would decrease. We should remember that 𝑘 itself is a free parameter that was fitted for each subject.

In the first model, on each trial, we assumed that subject updated the likelihood function with respect to 𝑡_𝑑_ as:

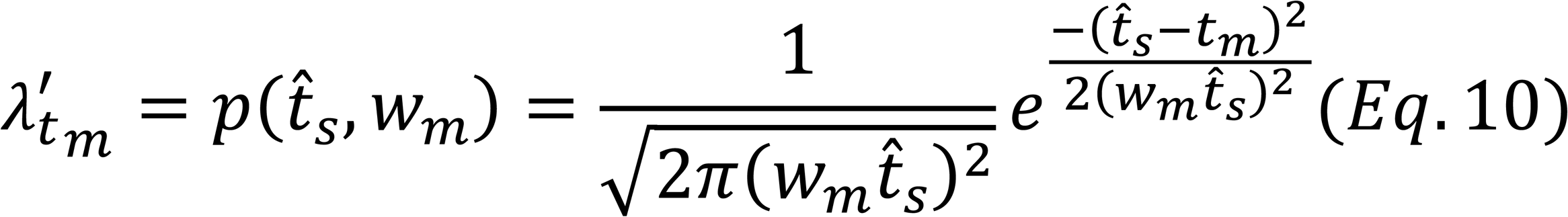

The direct relationship between the reproduced time and the sample time that was presented to the subject is explained as:

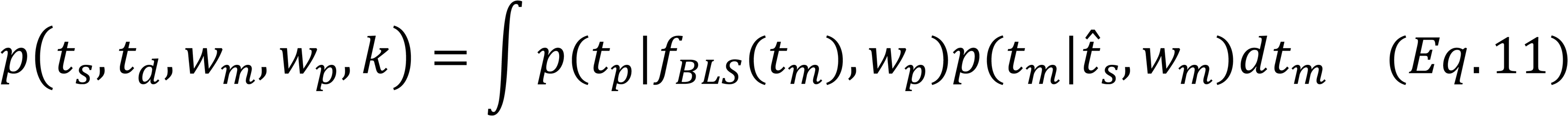

We named this model ‘Likelihood Model’.

For the second model, we applied the same logic of distractor to the value of 𝑡_𝑒_. The 𝑡_𝑒_ in the third step is replaced with 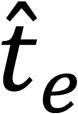 as:

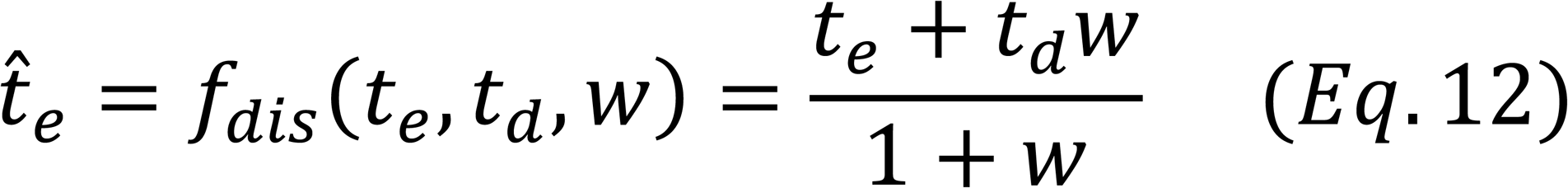

And the final equation of this model as:

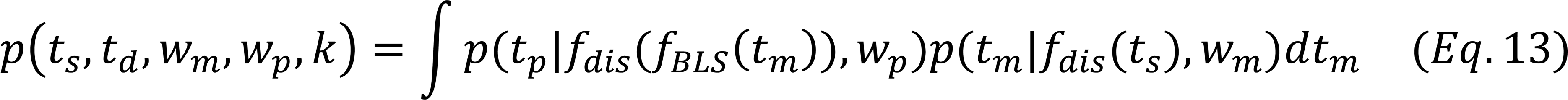

We named this model as ‘Motor Model’.

Delay interval can also affect time perception via updating the prior, similar to what was previously applied to sequential effects on timing [20]. To address this issue, we used an updating mechanism in which subject added a normal distribution (mean = 𝑡_𝑑_, sd = 𝑡_𝑑_*𝑤_𝑝_) to the initial prior. So, the updated prior for each trial computed as:

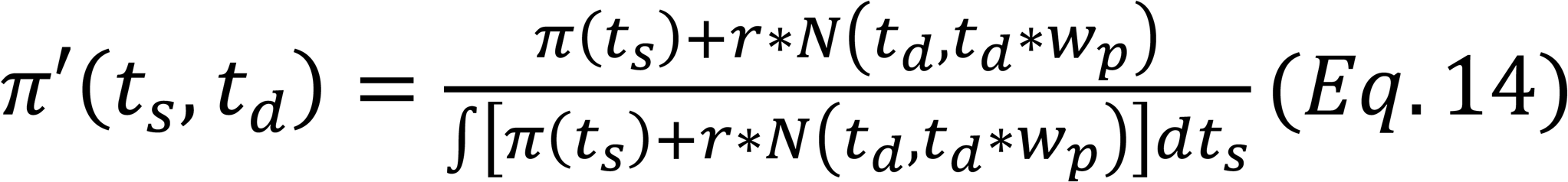

𝜋′(𝑡_𝑠_, 𝑡_𝑑_) is the updated prior and 𝑟 is the learning rate, so BLS function now incorporated the new prior to estimate the 𝑡_𝑒_ and the BLS formulation is:

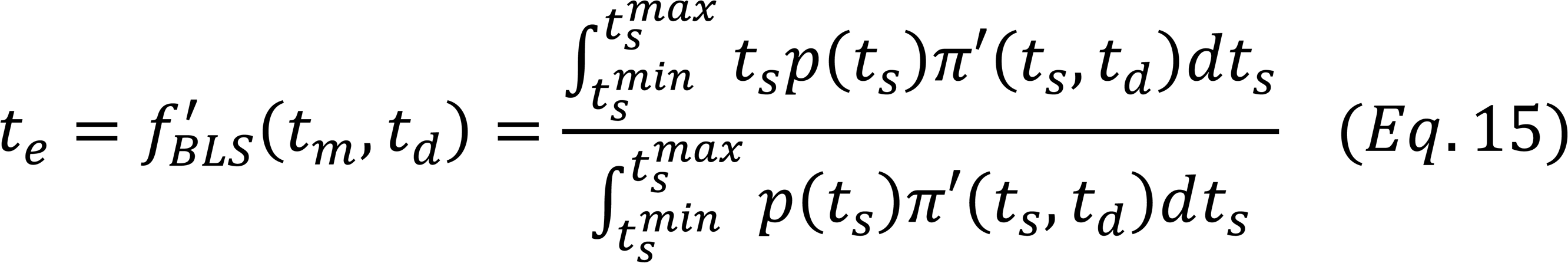

The final relationship between 𝑡_𝑠_ and 𝑡_𝑝_ is:

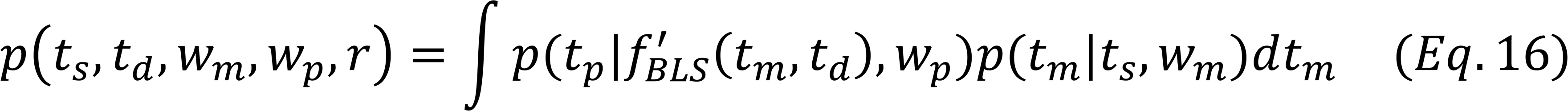

We called this third modeling the ‘Prior Model’.

We maximized the likelihood of model parameters 𝑤_𝑚_, 𝑤_𝑝_, and 𝑘 for the ‘Likelihood Model’ and ‘Motor Model’, and 𝑤_𝑚_, 𝑤_𝑝_, and 𝑟 for the ‘Prior Model’, across all combination of 𝑡_𝑠_, 𝑡_𝑝_, and 𝑡_𝑑_ for reproduction.

To compare the three fitted models, we first calculated Bayesian Information Criterion (BIC) as:

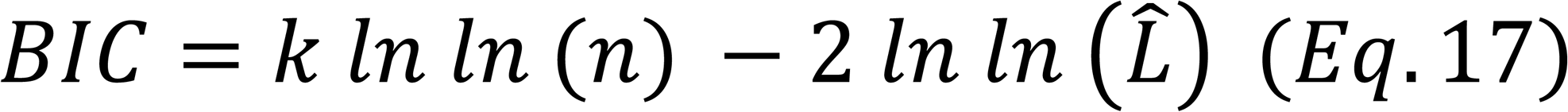

Which the 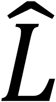 is the maximized value of the likelihood function of the model, 𝑛 is the number of observations (in our case the number of included trials for the reproduction task) and 𝑘 is the number of free parameters that are used in each model. We then calculated ΔBIC between the three models. Model with the smallest BIC would be the model with the best explanatory power (among the models that we studied). We use a Wilcoxon test to compare the median of the ΔBIC relative to zero to investigate whether it was significantly different from zero or not.

The modeling we explained so far, was applied to the reproduction task. For the discrimination task, we simulated data based on the cumulative Gaussian psychometric function fitted to the data (500 for each 𝑡_𝑠_ , 𝑡_𝑝_ combination, 4500 in total for each subject) as previously reported^22^.

## Results

In the discrimination task, the fraction of the trials that subjects reported that the second interval was longer than the first interval are plotted for 400 (Fig. 2A), 600 (Fig. 2B), and 1000ms (Fig. 2C) sample intervals, separately for 500 (green), 900 (blue), and 1700 ms (red) delay intervals. We found that as the sample interval increased, the sensitivity of time perception decreased, as expected according to Weber’s law, as can be appreciated by decreasing the slope of the fitted psychometric function as the sample interval increased. We then transformed the data based on the mean and standard deviation of the fitted cumulative Gaussian functions to the format in Fig. 2B, to compare data from discrimination and reproduction tasks directly. We also observed the classic Vierdordt’s effect of overestimation of shorter, and underestimation of longer durations in the time discrimination (Fig. 2E). Interestingly, it seems that the delay duration had no effect on time perception in the discrimination task. In the reproduction task (Fig. 2D), we observed Vierordt’s effect and Weber’s law, as expected. But in contrast to the discrimination task, the perceived time in the reproduction task was affected by the delay interval: as the delay interval increased, participants overestimated the sample interval.

**Figure 2.**
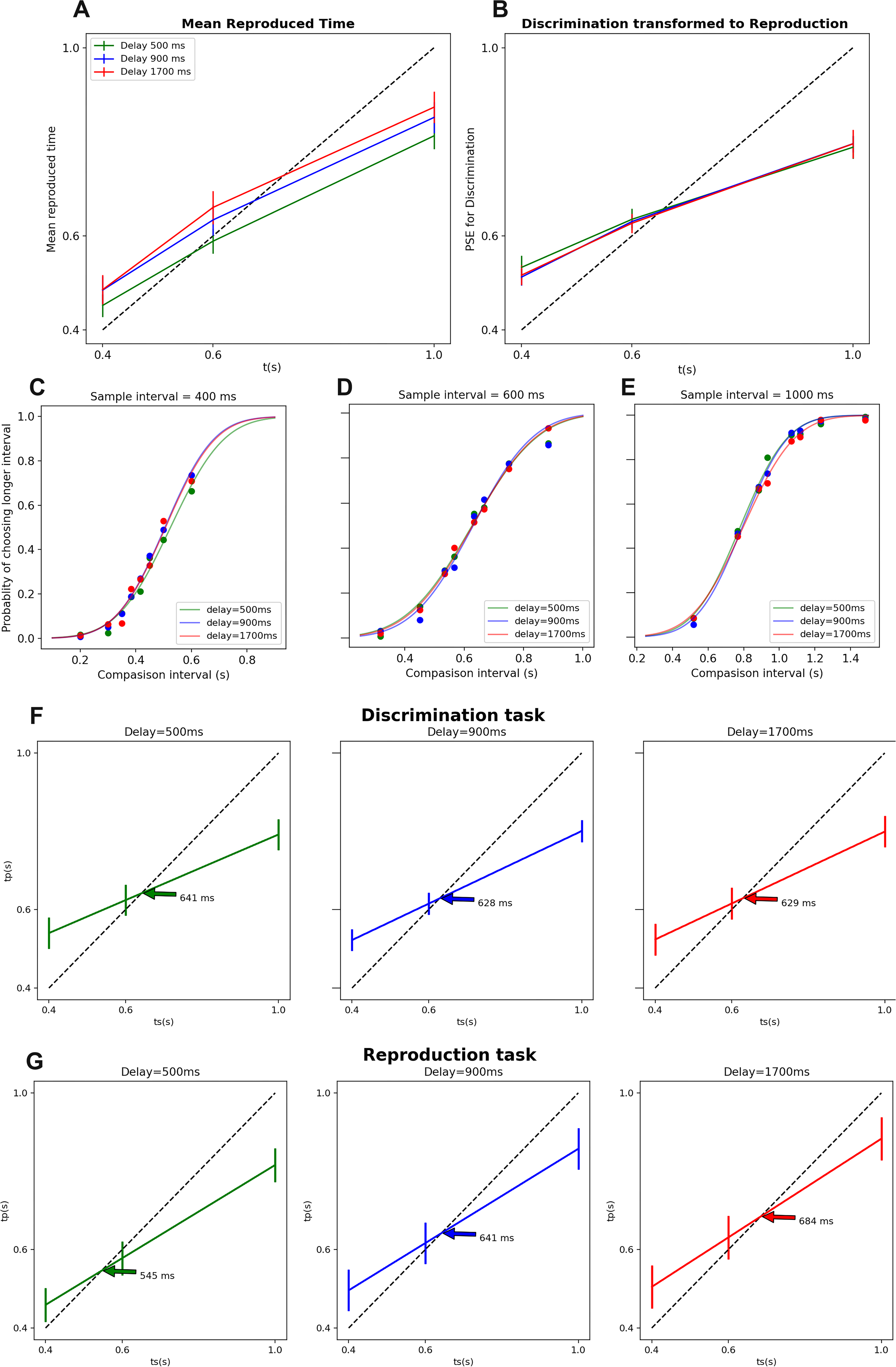
A, B and C, the proportion of long responses for different test interval duration and fitted psychometric function for 400, 600, and 1000ms delay interval, respectively. D, The mean of reproduced time as a function of sample interval in the reproduction task. E, The point of subjective equality (PSE) of perceived interval as function of sample interval in the discrimination task. F, The indifference points for the discrimination task for different delay intervals. Arrows in each subplot represent the indifference point. G, The indifference points for the reproduction task for different delay intervals. Arrows in each subplot represent the indifference point. In F and G error bars represent ± one standard error of the intercept.

In order to test the effects of the sample and delay intervals on the accuracy (i.e., mean) of the perceived time, we applied a three-way repeated measure ANOVA (Factor 1 = task: two levels; Factor 2 = sample interval: three levels; Factor 3 = delay interval: three levels). We found that while ‘task’ type (F = 0.12, df = 1, 0 = 0.73) or delay interval (F = 2.60, df = 3, p = 0.09) did not have any significant effects on the accuracy of time perception, sample interval had a significant effect (F = 186.89, df = 2, p < 0.0001). The interaction between sample interval × delay interval did not reach significant level (F = 0.71, df = 4, p = 0.58), but the interaction between task type × sample interval (F = 5.72, df = 2, p < 0.008) and task type × delay interval (F = 9.53, df = 2, p < 0.0008) reached significant level. The interaction between task type × sample interval × delay interval was not significant (F = 0.99, df = 4, p = 0.41).

Given the significant interaction between task type and other factors, we run a two-way RM ANOVA on the accuracy of time perception for discrimination and reproduction tasks, separately. In the discrimination task, while sample interval had a significant effect (F = 85.23, df = 2, p < 0.0001), delay interval did not (F = 0.03, df = 2, p = 0.96) on the accuracy. The interaction between sample interval × delay interval (F = 0.60, df = 4, p = 0.66) did not reach a significant level either. On the other hand, in the reproduction task, the effect of both sample interval (F = 144.29, df = 2, p < 0.0001) and delay interval (F = 20.57, df = 2, p < 0.0001) was significant. Their interaction, sample interval × delay interval, was not significant (F = 1.17, df = 4, p = 0.33). This result showed that delay duration affected the accuracy of time perception in the reproduction task (i.e., motor timing) but not in the discrimination task (perceptual timing).

Furthermore, we showed that the indifference point, the time of veridical judgment, in the discrimination task, was not different between delay durations: 636, 633, and 643ms in 500, 900 and 1700 ms delay durations (Fig. 2F, One-way ANOVA: F = 0.16, df = 2, p = 0.84). In the reproduction task we found that the indifference point increased from 571ms in 500ms delay to 713ms in 900ms delay and to 766ms in 1700 ms delay (Fig. 2G, One-way ANOVA: F = 10.02, df = 2, p < 0.006). The pairwise comparison was significant between all pairs: 500 vs. 900 (Signed Rank test, Z = 2.66, p < 0.003), 900 vs. 1700 (Z = 1.85, p < 0.03), and 500 vs. 1700 ms (Z = 2.79, p < 0.002).

To investigate the effect of the delay duration on the variability of perceived time, we applied a three-way RM ANOVA on standard deviation for each sample interval (3 levels), delay duration (three levels) and task (two levels). We found while task (F = 7.71, df = 1, p < 0.01) and sample interval (F = 13.18, df = 2, p < 0.0001) had a significant effect on the variability of time perception, delay interval did not (F = 1.88, df = 2, p = 0.17). The interaction between delay interval × task type (F = 1.46, df = 2, p = 0.24), between delay interval × sample interval (F = 1.09, df = 4, p = 0.36), between task × sample interval (F = 1.90, df = 2, p = 0.16), or between task × sample interval × sample interval (F = 0.17, df = 4, p = 0.94) did not reach significant level. Comparing these data with the results from effects of delay interval on the accuracy of time perception, we found that while delay interval led to overestimation, it did not have any effects on the variability of time perception.

Previous reports showed that in a reproduction task, preceding trials has a significant effect on the perception of time on the current trial ^16,17^. We were interested to see whether the delay interval, which comes after the sample interval in our task, had a similar effect on the current trial. To examine the effect of delay interval on sample interval, we calculated a deviation index. Deviation index is the difference between the reproduced time on a given trial and the subject’s mean reproduced time, regardless of delay duration, calculated separately for discrimination and reproduction tasks. The effect of delay interval on time perception in the reproduction task is shown by the change in the deviation index as a function of the difference between delay interval and sample interval (Fig. 3A). This deviation effect peaked when the delay interval was larger than the sample interval by approximately 900ms and decreased for larger differences. This interaction between the deviation index and the difference of the delay and sample interval was captured by derivative of Gaussian curve (r^2^ = 0.33), and also a linear fit (y = kx, r^2^ = 0.28) but not with the assumption that the deviation index was not dependent on the difference between sample and delay intervals (y = 0, r^2^ = -0.004). We applied the deviation index analysis to the data from the discrimination task as well (Fig. 3B). In line with the data presented in Fig. 2, we found that delay interval did not have any effects on the deviation index in the discrimination task (r^2^ = 0.001, 0.001, 0, for DoG, linear, and y = 0, respectively).

**Figure 3.**
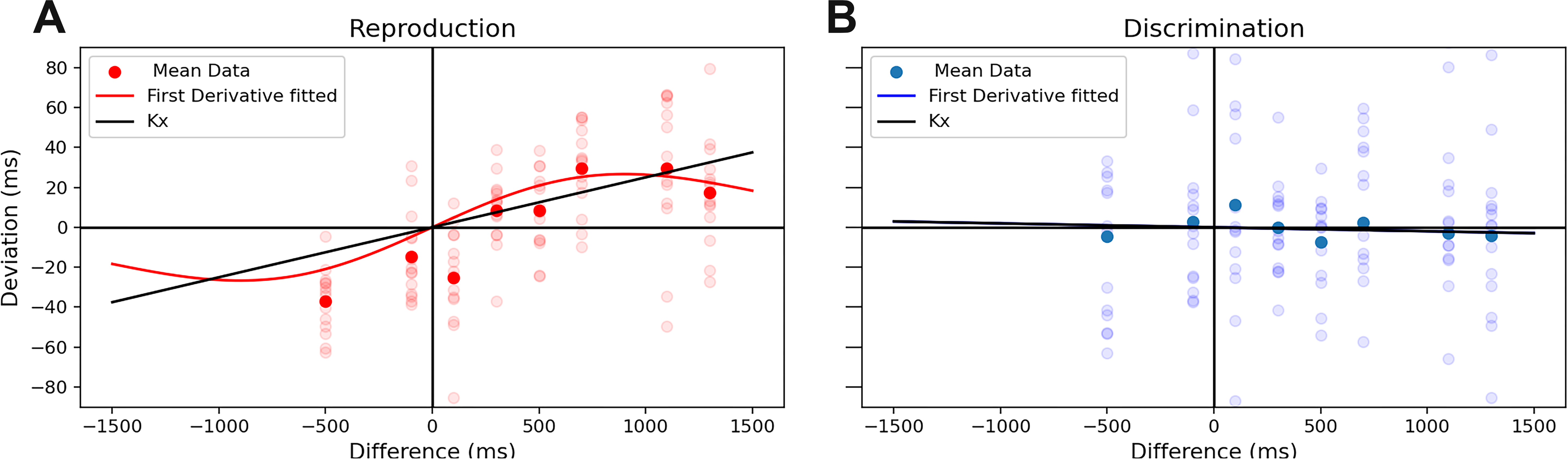
Effect of the delay interval on the deviation index in the reproduction (A) and discrimination (B) tasks. A: The mean of the deviation index as a function of difference between the delay interval and sample interval and fitted Derivative of Gaussian in reproduction task. Pale dots represent each subject’s data, and dark dots represent the average data. The red line represents the fitted derivative of the Gaussian (DoG) fitted to the population data. The black line represents the linear fit to the population data. B: The mean of deviation index as a function of difference between delay interval and sample interval and fitted Derivative of Gaussian in the reproduction task. Pale dots represent each subject’s data, and dark dots represent the average data. The blue line represents DoG fitted to the population data. The black line represents the linear fit to the population data.

So far, we showed that in the reproduction task, subjects overestimated time as a function of the delay duration. This could be explained by the fact that subjects were incorporating the delay interval into their timing process. This incorporation can happen at different levels. To tease apart underlying mechanisms quantitatively, we implemented models that took the delay interval into account, at three different stages of a classic observer model of time perception.

In the classic version of the model subjects combined the information from likelihood (Fig. 4A) with prior to come up with a posterior distribution (Fig. 4B) and then map the mean of the posterior to the reproduced interval with Gaussian distribution (Fig. 4C). To investigate the possible source for our findings we used three modified versions of the Bayesian observer model. In the first version of the model (Likelihood Model), the delay duration altered the mean and standard deviation of the *measurement* step with a weight of ’w’, determined by the leak factor (k). This incorporation changes mean and standard deviation of likelihood function (from solid to dotted line in Fig. 4D). In the second implementation, Prior Model, the delay interval alters the prior which is determined by the distribution of the sample interval (𝑡_𝑠_) on each trial (Fig. 4E, from green shade to gray). In this model the prior that is incorporated in the estimation step updates on each trial. Observer adds a normal distribution with mean of 𝑡_𝑑_ and standard deviation of 𝑡_𝑑_𝑤_𝑝_ to uniform prior distribution with the learning rate (𝑟). In the third version of the model (Motor Model), we incorporated the delay interval to the production stage of the Observer model (Fig. 4F, from solid blue to dashed). In this case, the mean and standard deviation of the distribution that maps estimated time to reproduced time were modified by the delay interval. We calculated the Bayesian information criterion (BIC) for each version of the model, fitted to each subject separately, to see which model explained our data best. We performed these stages for the motor timing task, since only in the motor timing task we observed that delay interval affected time perception.

**Figure 4.**
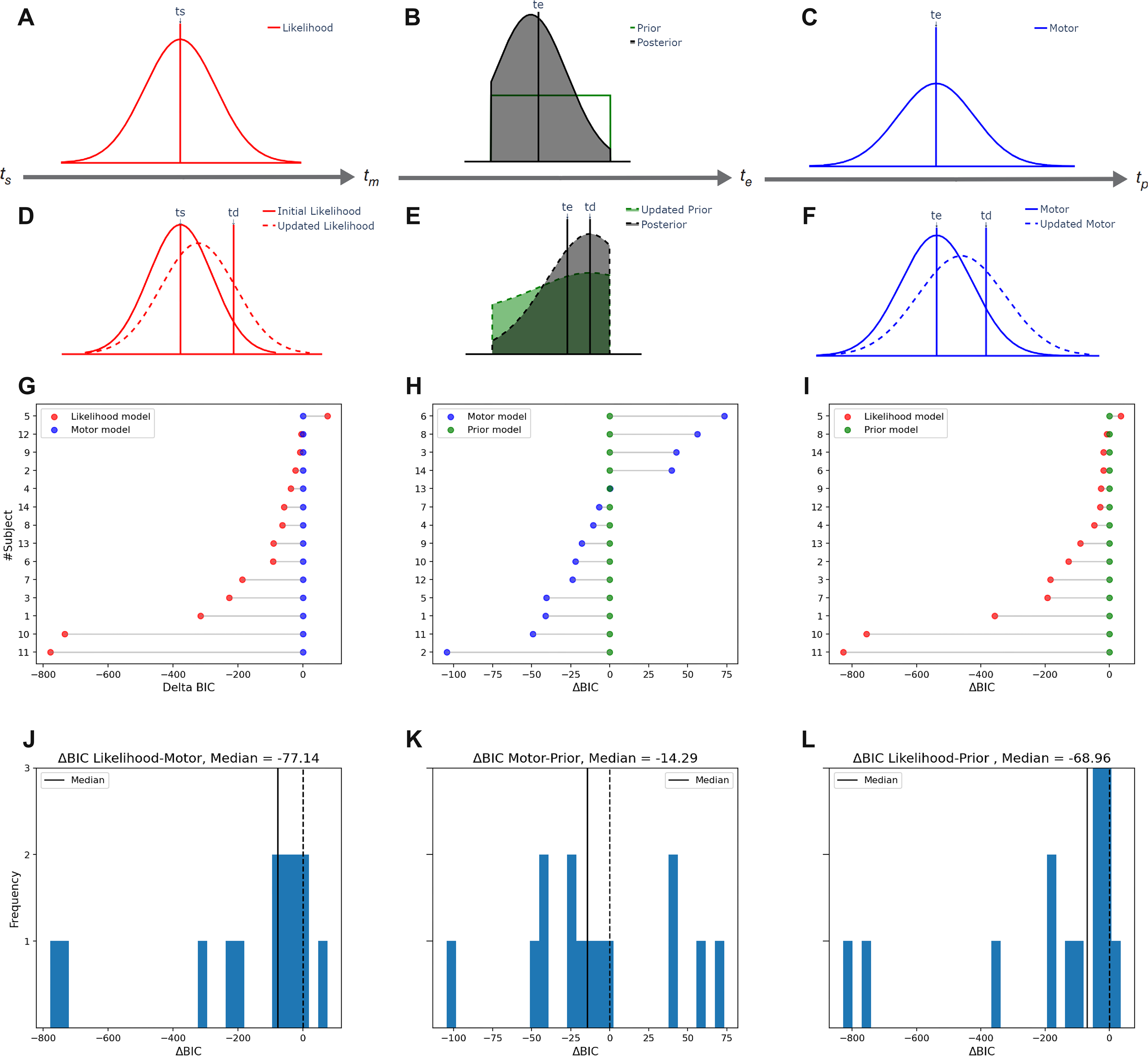
The three components of the classic Bayesian observer model: likelihood (A), incorporating prior (B) and motor component (C). The vertical line represents the sample duration on a given trial. The Gaussian distribution represents the likelihood with the mean of 𝑡_𝑠_ and standard deviation (SD) of 𝑡_𝑠_ × 𝑤_𝑚_. B, the green line represents the prior that was used in our study. The gray shaded area represents the posterior. The vertical line represents 𝑡_𝑒_ which is the average of the posterior distribution. C, the vertical line represents 𝑡_𝑒_ which is the same as 𝑡_𝑒_ in B. The Gaussian distribution represent the mapping function from 𝑡_𝑒_ to 𝑡_𝑝_ with the SD of 𝑡_𝑒_ × 𝑤_𝑝_. Modification applied to the Bayesian observer model in the likelihood function (D), prior distribution (E) and motor noise (F), which was performed independently and separately. D: The delay interval (𝑡_𝑑_) updates the likelihood of the sample interval (𝑡_𝑠_), from the solid, to the dotted red line, with the mean of 𝑡^_𝑠_ and the SD of 𝑡^_𝑠_ × 𝑤_𝑚_. E: In the Prior model, delay interval updates the prior and consequently the posterior. The updated prior (green shade) is the sum of the old prior with normal distribution centered on the 𝑡_𝑑_and SD of 𝑡_𝑑_ × 𝑤_𝑝_. The gray shade represents the consequently updated posterior. F: in the Motor model, the delay interval shifted the mean of production distribution from 𝑡_𝑒_ (with SD of 𝑡_𝑒_ × 𝑤_𝑝_, solid blue line) to 𝑡^_𝑒_ (with the SD of 𝑡^_𝑒_ × 𝑤_𝑝_, dotted blue line). G: ΔBIC between the Likelihood model and Motor model for each subject. Number on the y axis is subjects’ id number. ΔBIC between likelihood model and Prior model (H) per subject, and between the Motor model and Prior model per subject (I). Distribution of ΔBIC between likelihood model and Motor noise model (J), between likelihood model and Prior model (K) and between Motor model and Prior model (L). In J, K, and L, the dotted vertical line represents the value of 0, and the solid vertical line represents the median of the ΔBIC distribution.

We found that in the motor timing task, adding delay interval to the measurement stage (Likelihood Model) out-performed the model that added the delay interval to the production stage (Motor Model) in 13 out of 14 subjects (Fig. 4G). The median of ΔBIC was 77.14 smaller in the Likelihood Model than in the Motor Model (Fig. 4J) and was significantly lower than zero (Wilcoxon test, p < 0.02). On the other hand, the Motor Model was a better fit than the Prior Model in 9 subjects (Fig. 4H). The median of ΔBIC was 14.28 in favor of the Motor Model relative to the Prior Model (Fig. 4K, not significantly different from zero, p = 0.22). We found that the Likelihood Model also outperformed the Prior Model in 13 subjects (Fig. 4I). The median of ΔBIC was 68.95 in favor of the likelihood Model (Fig. 4L, significantly lower than zero, p < 0.03).

To be comprehensive, we applied these models to the data from the discrimination task. We found that the Likelihood model was a better fit than the Motor model in 8 subjects. Although the median of ΔBIC was 40.88 smaller in the Likelihood model than in the Motor model, this difference was not significantly different from zero (Figure S1A, p = 0.29). The Motor model was a better fit than the Prior model in 4 subjects, with median ΔBIC of 23.09, in favor of the Prior model (Figure S1B, p = 0.15). The likelihood model outperformed the prior model in 6 subjects with median ΔBIC 42.48, in favor of the Likelihood model (Figure S1C, p = 0.76)

Given that the Likelihood model emerged as the best-performing model in the reproduction task, and no model came on the top in the discrimination task, we compared the fitted parameter 𝑘, which represents the magnitude of integration of delay interval in sample interval, between the two tasks. We found that was significantly higher in the reproduction task (median = 0.256) than in the discrimination task (median = 0.002, Wilcoxon, p < 0.002, Figure 5).

**Figure 5.**
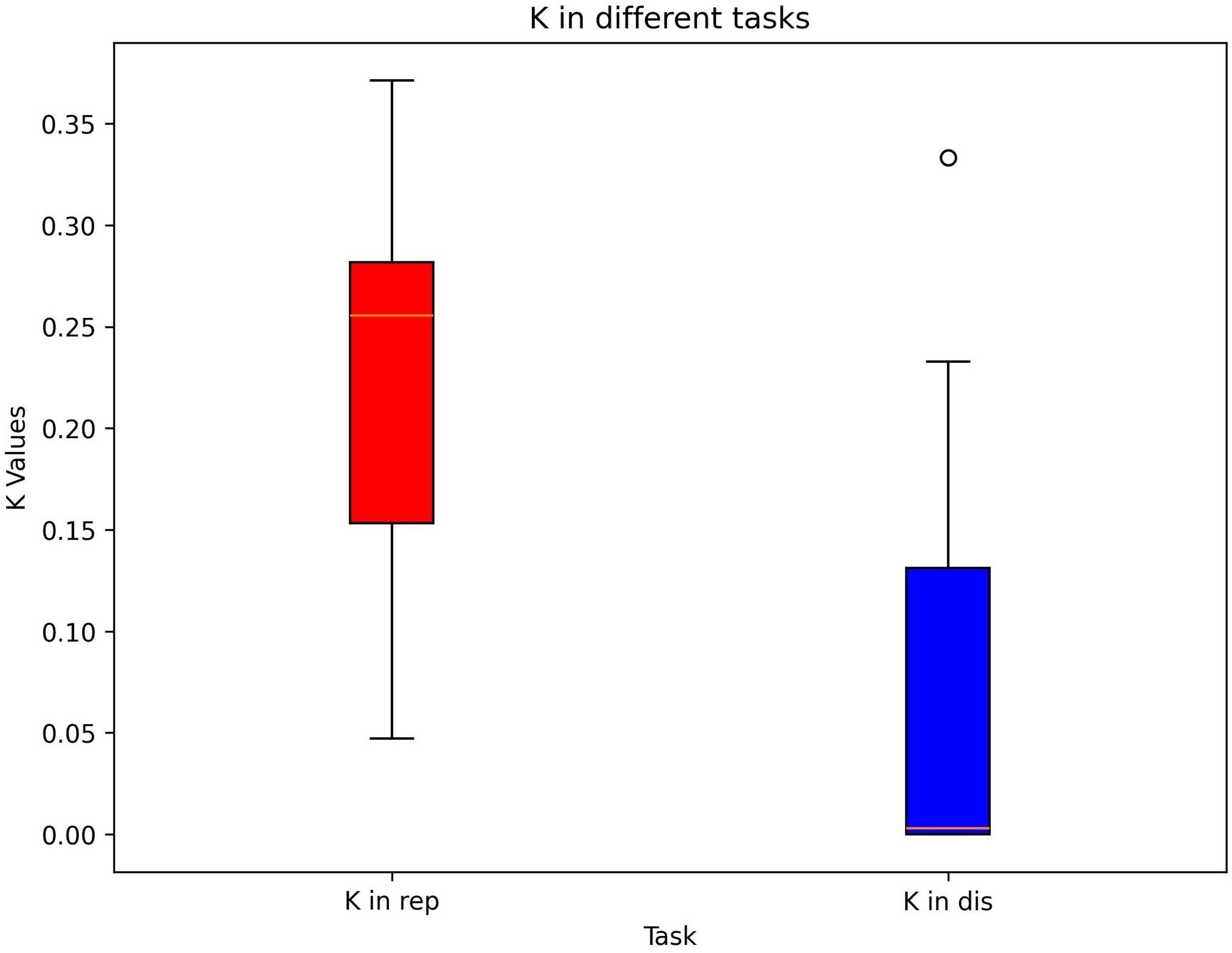
Comparison of the fitted parameter 𝑘 between the reproduction and discrimination tasks. The parameter 𝑘, representing the magnitude of integration between delay interval and sample interval, is significantly higher in the reproduction task (p < 0.002). The central box represents the interquartile range (IQR). The lower and upper edges correspond to the first (Q1) and third quartiles (Q3). The line inside the box indicates the median. The whiskers extend to the smallest and largest values within 1.5 times the IQR from Q1 and Q3, respectively. Data point outside this range is plotted as individual outlier.

## Discussion

We demonstrated that in a working memory task, increasing the delay interval, after a sample interval, led to an overestimation of the sample interval in a time reproduction task but not in the time discrimination task. More specifically, the perception of the sample interval was attracted towards the delay interval.We did not observe any changes in performance in the time discrimination task across different delay intervals between the first (sample) and the second (test) intervals. The effect of the delay interval on time perception in the reproduction task, can be interpreted not only as decreasing the fidelity of the representation of stimulus held in working memory, but also as biasing that representation towards the delay interval duration. We also found that this attraction towards the delay interval in the reproduction task can be better explained by updating the likelihood function rather than updating the prior or changes in motor processes.

Previous studies have shown that factors not relevant to the task, such as delay in our study, have an effect on the perception of time ^27^. One such study observed that the impact of inter-trial intervals on duration judgment is similar to that of delays ^27^.In another study, Heron et al., demonstrated that the presentation of auditory white noise before the sample duration as an adaptor could distract the perception of the subsequent sample duration. They showed that presentation of adaptor stimulus before a time discrimination task could cause attraction of points of subjective equality toward adaptor duration nonlinearly^28^.

There are different hypotheses about the underlying mechanism for this phenomenon, including the multi-channel process of time^29^. According to this theory, there is a specific population of neurons that each neuron encodes for a narrow range of duration, with overlapping tuning curves. The subsequent activation of different populations of neurons could cause distraction in the perceived duration^29^.

The observed overestimation of time in our reproduction task as a function of the delay interval has been reported before^30^. Fortin et al., hypothesized that this overestimation of sample duration in working memory tasks is due to fewer pulses accumulated in the pacemaker-accumulator model of time perception^31,32^. Based on these findings, we expected to observe that delay duration affects both motor and perceptual timing tasks, which is not what we found. We hypothesize that the underlying mechanism for this difference can be found in the two similar but slightly different aspects of memory: working memory and short-term retention of information. Tasks in which the subject is forced to perform a yes/no recognition instead of actively reproducing the stimulus held in memory are referred to as short-term retention tasks (like discrimination task in our study)^33^. On the other hand, tasks requiring active manipulation of the content of memory engage working memory. Different studies have shown that the circuits involved in each of these aspects of memory are distinct^34^. Postle et al., have shown that delayed recognition of the ordinal position of five randomly ordered letters in humans remains unaffected by repetitive transcranial magnetic stimulation (rTMS) of the mid-dorsolateral prefrontal cortex (dlPFC) during the delay period. However, impairment is observed with rTMS to the parietal cortex. On the other hand, rTMS of the dlPFC does affect performance when subjects are required to reorder these letters into alphabetical order^35^. So, our findings also show that increasing the delay duration affects the task which needs working memory (i.e., time reproduction) but not the task which only relies on the short-term retention of information (i.e., discrimination).

A previous study that used a delayed-response task to investigate the reproduction of time has reported different results from ours. Ueda et al., found that an increase in the delay duration caused an increase in the central bias of the reproduced time^36^. In their study, subjects overestimated the short duration more and underestimated the long duration also more as a function of the delay interval. Their data are not compatible with our finding that an increase in delay duration causes an increase in overestimation for both short and long durations. This difference in our data from theirs may be due to the difference in the time range between studies (0.4 to 1s in our case vs. 1 to 8s in theirs). Another possibility might be the difference in the fixation requirement in our design. In our task, subjects had to fixate on the center of the screen during the delay interval. We think that by maintaining fixation, our subjects were more aware of the passage of time compared to when they were free to look around. So, in our tasks, subjects might incorporate the delay duration into the sample duration. This can explain the discrepancy between their results and our data.

There is also another study which tackled the effect of working memory on time reproduction with a different task design^37^. In a n-back design, Gumus and Balci used a series of sample intervals, followed by an instruction to the subject to reproduce the nth sample interval. They demonstrated as the duration between the sample interval and time reproduction increased, the subject overestimated the sample interval. We believe that this effect is another manifestation of the effect observed in our data because the increase in n could be equivalent to an increase in the delay duration in our task. The similarity between the results of these two task designs shows that the overestimation due to an increase in working memory load is robust across different tasks.

To have a better understanding of how subjects incorporated the delay interval into their perception of time in the reproduction task, we used modified versions of the Bayesian observer model, introduced before^25^. It has been shown that the central tendency toward the mean of the prior distribution and scalar variability of time perception could be explained by Bayesian least squares estimator. Wang et al., investigated the serial dependence in time modality and showed that the preceding trial could affect the perception of the current trial by updating the prior^18^.

Contrary to their result, we found that the model which updates likelihood better explains our data in the reproduction task. We assume that this is due to the temporal organization of the delay interval in our task which is presented *after* the sample interval. We found that the prior is quite robust to the changes and manipulation which occur after the presentation of the stimulus, but at the same time, the likelihood function is prone to changes in temporal context (in our case, is presented after the stimulus). We also found that the likelihood model outperformed the model that changes the motor noise as a function of the delay interval duration.

In the discrimination task, none of the models outperformed other models, which is line with our empirical observation. Comparing the magnitude of integration of delay interval in sample interval, 𝑘, between two tasks, further showed that subjects integrated the delay interval in their estimation of time in the reproduction task, but not in the discrimination task. Given that none of the models surpassed the others in the discrimination task (perceptual timing task), we conclude that the integration of the delay duration with the sample duration (i.e., likelihood) is limited to the reproduction task (motor timing task).

In this study, we implemented the updating rule for likelihood in a way that it updates the mean and standard deviation of the likelihood function. There are also other implementations of likelihood update that could explain the biases away from the prior distribution. In this line, Wei and Stocker showed that the asymmetric likelihood duration could explain the repulsion from the mean of prior distribution^38^. We assume that these two implementations could change the posterior in a way that leads to an increase in the density of posterior, in the areas away from the mean of the prior. The stability of the prior and the trial-by-trial updates in the likelihood function show that subjects have both stability in their perception and sensitivity to changes in temporal context, which lead to a better estimation of a sensory stimulus^39^.

## Data availability

Data from subjects and corresponding codes for data analysis and modeling will be made available online upon publication of the manuscript.

## Acknowledgment

We would like to thank Ahmad Pourmohammadi and Sepehr Sima for their helps in model implementation and comments on the previous version of the manuscript.

---

**Figure S1.**
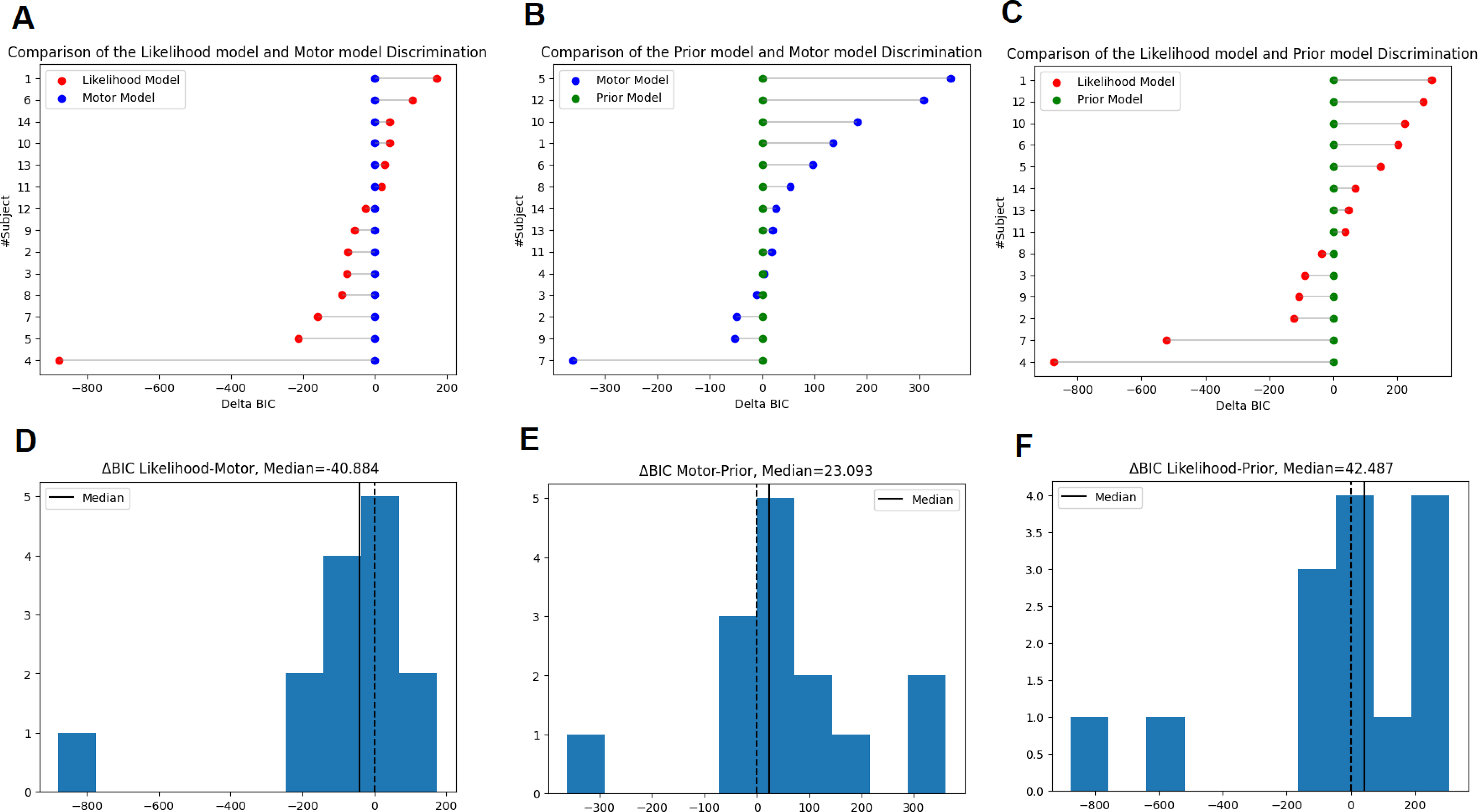
**Top A**: ΔBIC between the Likelihood model and Motor model for each subject. Number on the y axis is subjects’ id number. **B**: ΔBIC between likelihood model and Prior model per subject. **C:** ΔBIC between the Motor model and Prior model per subject. **Bottom:** Distribution of ΔBIC between each two models comparison.

